# Olfactory receptors are sensitive to molecular volume of odorants

**DOI:** 10.1101/013516

**Authors:** Majid Saberi, Hamed Seyed-allaei

## Abstract

To study olfaction, first we should know which physical or chemical properties of odorant molecules determine the response of olfactory receptor neurons, and then we should study the effect of those properties on the combinatorial encoding in olfactory system.

In this work we show that the response of an olfactory receptor neuron in Drosophila depends on molecular volume of an odorant; The molecular volume determines the upper limits of the neural response, while the actual neural response may depend on other properties of the molecules. Each olfactory receptor prefers a particular volume, with some degree of flexibility. These two parameters predict the volume and flexibility of the binding-pocket of the olfactory receptors, which are the targets of structural biology studies.

At the end we argue that the molecular volume can affects the quality of perceived smell of an odorant via the combinatorial encoding, molecular volume may mask other underlying relations between properties of molecules and neural responses and we suggest a way to improve the selection of odorants in further experimental studies.

## 1 Introduction

Survival of many species depends on their olfactory system. They use it to search for food, avoid poison, escape from danger, find mate, and bind to their offspring. An olfactory system detects volatile chemicals in the surrounding, encodes the results and transmit them to limbic system and cortex.

The front end of the olfactory system are olfactory receptor neurons. Each neuron expresses only one kind of olfactory receptor (in insects they are co-expressed with Orco [1]), neurons of the same type converge into the same glomeruli of the olfactory bulb (or antenatal lobe in insects), so that each glomerulus of olfactory bulb receives an amplified signal from only one type of olfactory receptor [2–10].

From neural recordings we know that the olfactory systems use a combinatorial code: an olfactory receptor can be triggered by different odorant molecules, and an odorant molecule can excite different olfactory receptors [11]. The combinatorial code helps the olfactory system to discriminates trillion odors [12]. However, it is not clear yet which properties of a molecule contribute to its smell, it is a topic of ongoing researches and there are many theories [13–24].

In this study, we investigate the relation between molecular volumes of odorants and the responses of olfactory receptor neurons. Our results suggest that molecular volume is a considerable factor, but not the only factor that determines the neural response of the olfactory receptor neurons.

The olfactory receptors are transmembrane proteins. In vertebrates, they are metabotropic receptors, they belong to the family of g-protein coupled receptor (GPCR), Linda B. Buck and Richard Axel won the Nobel Prize in Physiology or Medicine, in 2004, for the discovery of this [25]. There are many similarities between the olfactory system of insects and vertebrates [26, 27], and it was assumed that insects use the same kind of signal transduction [28, 29], but recently, it has been argued that the olfactory receptors in insects are inotropic [30–33], their topology is different from vertebrates [34, 35], and they function in presence of another common receptor, called Orco [1].

Regardless of the signal transduction, all olfactory receptor have the same function, they have a binding-pocket (also known as binding-cavity and binding-site), where the ligands (odorants) bind to. This binding activates the receptors and the activated receptor changes the potential of the cell, directly (inotropic) or indirectly (metabotropic).

The amount of change in the membrane potential of a olfactory receptor neuron depends on the number of activated olfactory receptor proteins and the time that they remain activated, which are determined by various physio-chemical properties of the ligand (odorant) and the receptor [13, 15, 19], but here we focus only on two of them: the volume and the flexibility of the binding-pocket. The molecular volume of a ligand should match the dimensions of the binding-pocket of the receptor, then it fits into the binding-pocket of the receptor and triggers the signal transduction. Any mismatch in the volumes will affect the neural responses (Fig. 1a), on the other hand the flexibility of the binding-pocket can compensate for the volume mismatch (Fig. 1b),

**Fig. 1.**
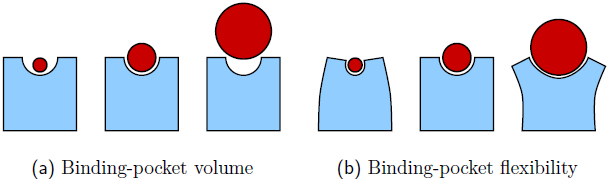
This figure shows different scenarios that may happen when an odorant molecule (ligand) binds to a receptor. Fig. 1a shows the effect of binding-pocket volume. From left to right, misfit because of small volume of molecule, perfect match and misfit because of large molecular volume. Fig. 1b demonstrates that the flexibility of a receptor may compensate for the volume mismatches. The red disks (dark grey in b&w) are odorant molecule, and the blue shapes (light grey in b&w) are olfactory receptor and binding-pocket.

We could know the volume and flexibility of the binding-pocket, if we knew its three dimensional structure. But this is not the case here, it is not easy to know the structure of integral proteins [36, 37], including olfactory receptor. It is the topic of ongoing researches, using various methods like Molecular Dynamic (MD) simulations, mutagenesis studies, heterologus expression studies, and homology modeling [38–46]. In this study, we use neural recording to predict the volume and flexibility of binding-pocket of olfactory receptors, *in-vivo*.

In this study we suggest a functional relation between molecular volume and the neural responses, we provide a methodology to estimate *chemical range* or *tuning function* of olfactory receptors, and then we predict the structural properties of the binding-pocket of olfactory receptor - the volume and the flexibility of binding-pocket. Our results may help to odorant selection of new experimental studies, may provide additional information about the structure of olfactory receptors to structural biologists, and may contribute to the study of olfactory coding.

To perform this study we use a public domain, well structured database – DoOR – that includes the neural responses of most olfactory receptors (OR) of Drosophila to many odorants [47]. This database has collected its data from many other sources [18, 20, 48–60].

## 2 Material and methods

We want to study the relation between neural responses and molecular volumes, so we need the respective data. We take the neural data of DoOR database [47] and we calculate molecular volume (supplemental file 3) using a computational chemistry software – VEGA ZZ [61]. We used GNU R to analyse the data [62].

DoOR database can be summarized in an *N* × *M* matrix. Its elements *r*_*nm*_, are the response of neuron *n* to odorant *m*. This matrix is normalized between 0 and 1 so we have 0 ≤ *r*_*nm*_ ≤ 1, where 1 is the strongest response. The only problem is that this matrix has some Not Available (NA) values, different neurons are excited by different set of odorants, so when summing over *m*, Σ_*m*_, we are calculating Σ_*m*:*r*_*nm*_≠NA_, but for simplicity, we use the former notation.

The response *r*_*nm*_ depends on the molecular volume of the odorant, *v*_*m*_, and other physio-chemical properties of the molecule *m*; We assume that we can separate the response *r*_*nm*_ into two terms:

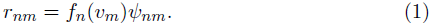

The first term, *f*_*n*_(*υ*_*m*_), depends only on the molecular volume of odorants.

The second term, *ψ*_*nm*_ include every other influential properties of molecules, but the molecular volume. Both terms are characteristic of each receptor, and they might vary from neuron to neuron. In fact, the first term, *f*_*n*_(*υ*), is the tuning curve of neuron *n* in respect to the molecular volumes, it can be approximated with a Gaussian function

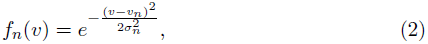

where, *υ*_*n*_ is the preferred molecular volume of receptor *n* and *σ*_*n*_ represents its flexibility. In this work we want to estimate *υ*_*n*_ and *σ*_*n*_. To do so, first we calculate the response weighted average of molecular volumes, 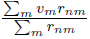 and then we use (1):

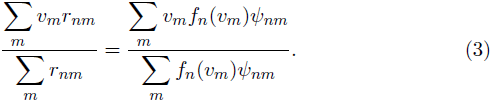

Here we can approximate Σ with ∫, which is common in statistical physics:

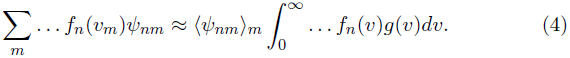

In which, 〈*ψ*_*nm*_〉_*m*_ denotes the average of *ψ*_*nm*_ over all *m* : *r*_*nm*_ ≠ NA. It can be moved out of the integral for it is independent of *ψ*. In the above equation, *g*(*υ*) is the density of states, *g*(*υ*)*dυ* indicates how many molecules have a molecular volume in the range of *υ* and *υ* + *dυ*. This function can be approximated by a Gaussian function, Fig.2,

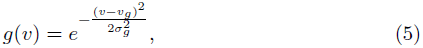

ideally, *g*(*υ*) should not depend on the neuron *n*, it is the property of ensemble of odorant molecules, not neurons. But here, we have many missing values (*r*_*nm*_ = *NA*), so we have to calculate *g*(*υ*) for each neuron separately; Therefore, *υ*_*gn*_ and *σ*_*gn*_ are the average and standard deviation of molecular volume while *r*_*nm*_ ≠ NA. Now we rewrite equation (3) using equation (4):

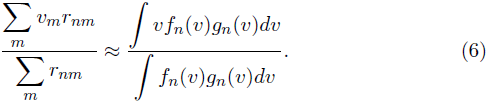

We replace the product of *f*_*n*_(*υ*) and *g*_*n*_(*υ*) in the above equation with *h*_*n*_(*υ*) = *f*_*n*_(*υ*)*g*_*n*_(*υ*), to make a simpler form

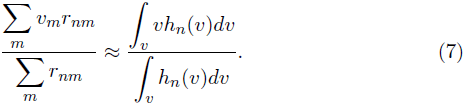

The function *h*_*n*_(*υ*) is a Gaussian function because it is the product of two Gaussian functions,

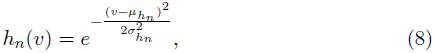

so the right hand side of equation 7 is nothing but *μ*_*h*_*n*__ and in a similar way, we can calculate *σ*_*h*_*n*__ from the neural data

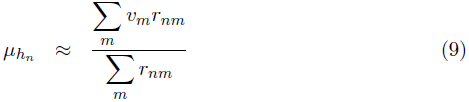

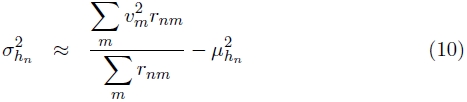

We knew the mean *υ*_*g*_*n*__ and standard deviation *σ*_*g*_*n*__ of *g*_*n*_(*υ*) from the molecular volumes of the ensembles of odorants. We just calculated the mean *μ*_*h*_*n*__ and standard deviation *σ*_*h*_*n*__ of *h*_*n*_(*υ*) from the neural data. Now calculating the mean *υ*_*n*_ and the standard deviation *σ*_*n*_ of *f*_*n*_(*υ*) is trivial, first we calculate *σ*_*n*_ from

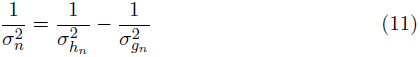

and then we calculate *υ*_*n*_:

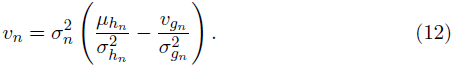

The calculated *υ*_*n*_ and *σ*_*n*_ are in supplemental file 1. The resulting *f*_*n*_(*υ*) are plotted over the actual data, for 32 receptors (Fig. 3a), in which the relative error of *υ*_*n*_ is lesser than 25% and *σ*_*n*_ < 80*Å*^3^, and for one receptor just magnify the details (Fig. 3b). Now we know the preferred volume *υ*_*n*_ of each receptor and also its flexibility *σ*_*n*_.

**Fig. 2.**
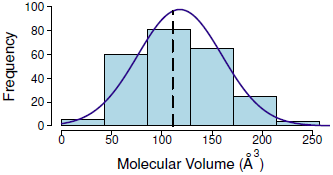
Density function of molecular volumes (*g*(*υ*)), considering all molecules of DoOR database. The actual density function of molecular of volumes in each experiment (*g*(*υ*)) might be slightly different because each experiment uses a different subset of molecules. The solid line is a Gaussian fit (Eq. 5) and the dashed line shows the median, which is slightly different from the mean.

**Fig. 3.**
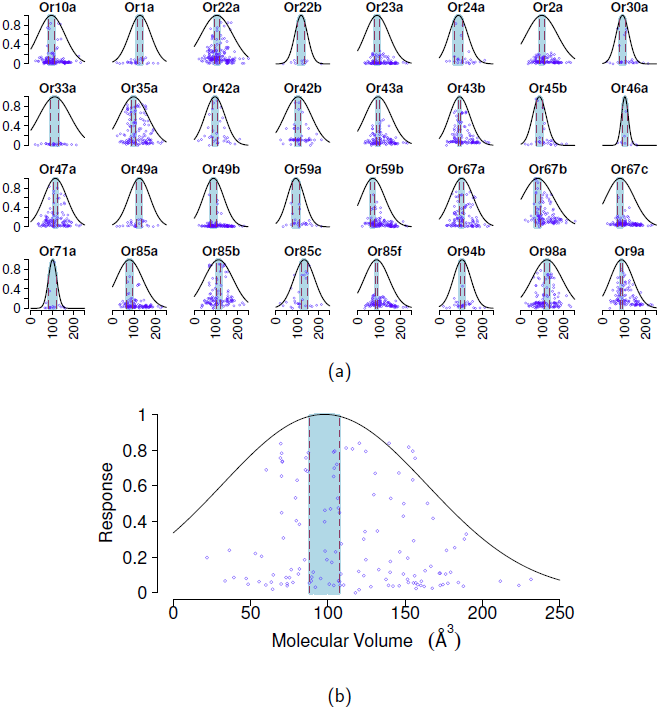
Response of olfactory receptors versus molecular volume of odorants (Circles), the fitted functions *f*_*n*_(*υ*) from Eq. 1 (solid lines), and the error bars of the mean of *f*_*n*_(*υ*) (red vertical lines), for 32 selected receptors (Fig. 3a) and for one selected receptor Or35a (Fig. 3b) just to magnify details.

## 3 Results and discussions

There are two main assumption in this work: First we assumed that the response of an olfactory receptor can be factorized into two terms, according to (1). Second, we assumed that the volume dependence factor *f*_*n*_(*υ*_*m*_) in (1) have a Gaussian form (Eq. 2). Considering the physics and chemistry behind the binding-process (Fig. 1), and the neural responses (Fig. 3), these assumptions are logical.

The function *f*_*n*_(*υ*) can be considered as the tuning curve of olfactory receptor n in response to molecular volume (Fig. 3). Each receptor has a preferred molecular volume *v*_*n*_ and shows some flexibility *σ*_*n*_. We calculated the parameters of *f*_*n*_(*υ*) for 32 receptors (Fig. 3). The calculated values, *v*_*n*_ and *σ*_*n*_ are in Fig. 4a and 4b respectively. Figure 4a demonstrate that the molecular volume preference of receptors are different. Figure 4b illustrate that the flexibility of receptors are also different.

**Fig. 4.**
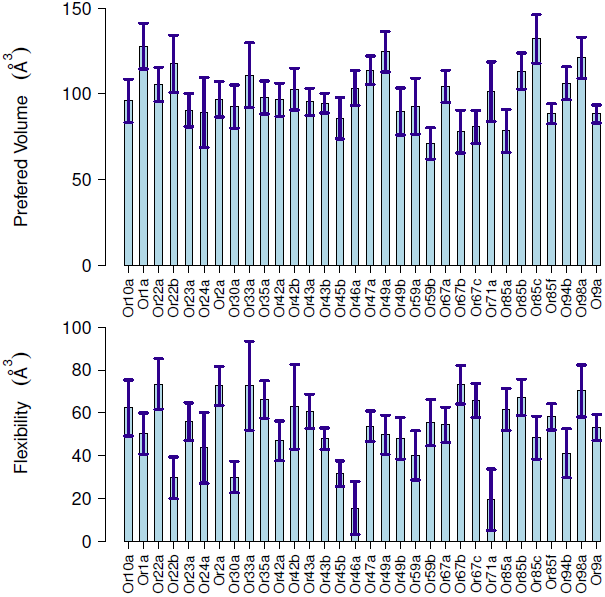
(a) The preferred volumes of 32 receptors (*υ*_*n*_), and their error bars. The error bars are calculated using Jack-Knife method. Some receptors prefer smaller molecules - like *Or59b*, *Or67b* and *Or85a*, but some other receptors prefer larger molecules - like *Or85c*, *Orla* and *Or49a*. (b) The flexibility of each receptor (*σ*_*n*_), the error bars are calculated using Jack-Knife method. Some receptors like *Or46a*, *Or71a* and *Or22b* are volume selective, but some other receptors like *Or22a*, *Or67b* and *Or33a* show flexibility and respond to broader range of molecular volumes.

This diversity is important in perceiving the quality of smells. In a hypothetical experiment, assume that every characteristic of odorant molecules are the same but their molecular volume. If all olfactory receptors had the same preferred volume and flexibility, any change in the molecular volume would change only the intensity of smell not its quality. But olfactory receptors have different preferred volumes and flexibilities, so any change in the molecular volume of an odorant results in a different combinatorial encoding which affects the quality of perceived smell as well. That may describe the difference in the smell of methanol, ethanol, propanol and butanol. Methanol smells pungent, ethanol smells pleasant and winy, propanol smells like ethanol while butanol is similar to ethanol with little banana like aroma. The molecular volume affects the combinatorial encoding.

Here we showed that the responses of olfactory receptor neurons are related to the molecular volume of odorants, apart from that, it is not clear which other features of molecules are measured by olfactory receptors. It is a topic of ongoing researches, there are many works that try to connect the physio-chemical properties of molecules to the evoked neural response or perceived smells. But the non-linear volume dependence (Eq. 1 and Eq. 2) may mask important relations between molecules and neural responses.

By considering the effect of molecular volume on the response of olfactory receptor neurons, one might discover more subtle dependence between other molecular features and neural responses, by studding *ψ*_*nm*_, which otherwise would be masked by this non-linear relation *f*_*n*_(*υ*).

We also predict some *in-vivo* structural aspects of the binding-pocket of olfactory receptors: the preferred volume of each receptor results from the volume of the binding-pocket, the flexibility of a receptor results from the rigidity or flexibility of the binding-pocket; These data add some constrains over the 3d structure of olfactory receptors, which may help the prediction and calculation of 3d structure of these proteins.

The method of this work can be combined with mutagenesis as well. Some genes of an olfactory receptor are mutated, then its response to a selection of molecules are measured and finally the preferred volume and flexibility are calculated. In this way we can understand which amino acids of the olfactory receptor contribute to the volume and flexibility of the binding-pocket, as well as affecting the function of the receptors.

Our finding can also save time and expenses of experiments by suggesting important odorants for every receptors. To study *ψ*_*nm*_ of a receptor, it is better to have many data points and those data points are better to be around the preferred volume of the receptor. But this is not the case in current data, for many receptors, most data points are on the tails of *f*_*n*_(*υ*), which is close to zero. We suggested the best selection of odorants for each of 32 studied receptors (see Venn diagram in Fig. 5 and supplemental file 2).

**Fig. 5.**
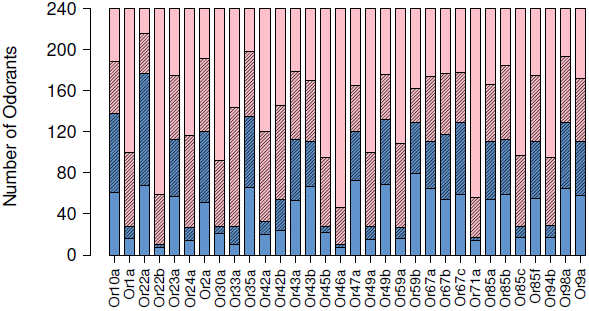
Venn diagram of DoOR database and our suggested important odorants of each receptor. The database includes 240 odorant molecules, only a fraction of them are used to study an olfactory receptor (blue shades) and data for the rest of odorants are not available (pink shades). The hatched area are odorants with molecular volume close to the preferred volume of each receptor (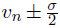). We consider them as important samples to study the molecular basis of olfaction, We already know the neural response of hatched blue area, but the hatched pink odorants could be the target of further experiments.

Although this work is on the data of Drosophila, we expect that the general principle and methodology of this work hold for vertebrates as well. But considering the similarities and dissimilarities between insects and vertebrate, this should be verified and more work are necessary.

## 4 Acknowledgments

We are especially grateful to B. N. Araabi, S. Aghvami and N. Doostani for the careful reading of the manuscript; and P. Carloni for the fruitful discussion.

